# Adult Drosophila aversion to caffeine requires a unique TrpA1 isoform and the PLC signaling cascade

**DOI:** 10.1101/2025.11.20.689420

**Authors:** Romane Milleville, Jade Gillas, Amanda Thuy La, Gérard Manière, Martine Berthelot-Grosjean, Yaël Grosjean, Julien Royet, C. Leopold Kurz

## Abstract

Taste in Drosophila melanogaster is crucial to survival, influencing feeding, mating, and egg-laying behaviors. Taste organs are located on various parts of the body, including the legs, proboscis, wings, and ovipositor. Taste neurons detect chemicals via receptors like GRs, IRs, and TRPs, with bitter and sweet tastes linked to specific neurons (*Gr66a+* and *Gr5a+*). Bitter substances such as caffeine activate neurons, resulting in rejection behavior. TrpA1 channels, associated with aversive responses, are involved in complex behaviors and could interact with taste receptors. Our results show that caffeine mixed with sucrose reduces proboscis extension in flies compared to sucrose alone, a response that requires only the TrpA1-E isoform out of the five possible ones. Furthermore, our data demonstrate that this avoidance requires TrpA1 and signaling via PLC and IP3-receptors in adult *Gr66a+* neurons.

## INTRODUCTION

The sense of taste plays a fundamental role in environmental assessment, allowing organisms to make crucial survival-related decisions. In *Drosophila melanogaster*, gustatory perception regulates key behaviors such as feeding (Dweck & Carlson, 2023; Thorne et al., 2004), mating (Miyamoto and Amrein, 2008), and oviposition (Chen and Amrein, 2017; Joseph et al., 2009; Montell, 2009). A central challenge in food selection lies in distinguishing caloric sources (Dus et al., 2011; Stafford et al., 2012) from toxic compounds (Delventhal et al., 2017) and evaluating mixtures of both (French et al., 2015).

The peripheral taste organs of *D. melanogaster* are distributed across several body parts, including the legs (Liman et al., 2014; Ling et al., 2014), wings (Raad et al., 2016), ovipositor (Yang et al., 2008), and the proboscis—a long appendage functionally analogous to the human tongue (Dweck and Carlson, 2023; Stocker, 1994). Taste neurons housed within bristles on these organs directly interact with chemical stimuli through dendrites that contact solid or liquid surfaces. Although individual neurons can detect multiple compounds, their behavioral output is typically consistent (Sung et al., 2017; Wang et al., 2004). This specificity arises from the co-expression of multiple receptor classes, including gustatory (GR), ionotropic (IR), and transient receptor potential (TRP) channels (Chen and Amrein, 2017; Freeman and Dahanukar, 2015; Koh et al., 2014).

Among the best-characterized gustatory neurons are those expressing the Gr66a receptor ( cells), which mediate bitter taste (Thorne et al., 2004; Wang et al., 2004), and Gr5a-expressing neurons (*Gr5a+* cells), which respond to sugars (Dahanukar et al., 2001; Ueno et al., 2001). Sweet compounds such as sucrose activate *Gr5a+* neurons, whereas bitter substances like caffeine stimulate neurons, where Gr66a serves as a co-receptor (Amrein and Thorne, 2005; Lee et al., 2009; Moon et al., 2006). Activation of these distinct neuronal populations leads to acceptance or rejection of food, respectively (Thorne et al., 2004).

Beyond gustation, neurons contribute to aversive responses to environmental stressors such as UV radiation through the TRP channel TrpA1 (Guntur et al., 2017; Rosenzweig et al., 2005). TrpA1 has been widely studied in *Drosophila*, where it mediates responses to temperature extremes (Gu et al., 2019), mechanical stimuli (Gong et al., 2023, 2022), and chemical irritants (Kim et al., 2010). While caffeine modulates TRPA1 activity in mammals (Nagatomo and Kubo, 2008), *Drosophila* TrpA1 does not appear to influence caffeine avoidance in binary choice assays or taste sensilla recordings (Kim et al., 2010). Nonetheless, TrpA1 may participate in more complex aversive mechanisms through interactions with gustatory receptors (Kwon et al., 2010).

Feeding behavior in *Drosophila* follows a sequence of sensory checkpoints—from olfactory detection to leg contact, proboscis engagement, ingestion, and consumption (Thoma et al., 2017). Given TrpA1’s role in detecting environmental threats, we investigated whether this channel mediates caffeine avoidance at specific stages of feeding. We focused on the proboscis and its associated feeding reflex, the Proboscis Extension Reflex (PER), in which flies extend the proboscis in response to palatable stimuli (Schwarz et al., 2017).

We found that adding caffeine to sucrose significantly reduced PER compared to sucrose alone. Of the five known *Drosophila* TrpA1 isoforms, only TrpA1-E was required for this effect. Caffeine-mediated suppression of PER depends on *TrpA1* expression in neurons and involves a signaling cascade including phospholipase C (PLC) and the inositol trisphosphate receptor (IP3R).

These findings reveal a more nuanced role for TrpA1 in taste aversion, extending its function beyond nociception to cooperative signaling with gustatory pathways during feeding-related decision-making.

## RESULTS

### Caffeine Elicits Aversive Responses in a Proboscis Extension Reflex Assay

Caffeine is widely recognized as an aversive compound in *Drosophila*, with behavioral quantification typically performed using two-choice assays (Lee et al., 2009) or the multiCAFE assay (Sellier et al., 2011). These paradigms are effective for assessing attraction or aversion to specific chemicals and for quantifying consumption when flies freely explore their environment in groups over periods ranging from minutes to hours.

To investigate whether TrpA1 is necessary for caffeine-induced aversion and to delineate the neuronal populations and molecular pathways involved, we focused on a reflexive feeding behavior: the Proboscis Extension Reflex (PER). PER is triggered when taste sensilla on the legs or proboscis contact a stimulus. A palatable substance induces immediate proboscis extension, whereas aversive compounds suppress it. In this study, we selectively stimulated the proboscis. To distinguish active aversion from simple disinterest, caffeine was tested in combination with 1 mM sucrose, which reliably induces PER (Video 1). A significant reduction in PER when caffeine is added to sucrose indicates an aversive response rather than a lack of interest.

Historically, two-choice arenas, multiCAFE assays, and electrophysiology have been the preferred methodologies for assessing responses to caffeine, with PER used primarily to validate its aversive properties (Mi et al., 2023). To establish the optimal caffeine concentration for our study, 5- to 7-day-old *w⁻*females were exposed to increasing caffeine concentrations mixed with 1 mM sucrose. Consistent with previous findings (Montanari et al., 2024), sucrose alone elicited PER in approximately 80% of flies (Fig. 1a; Video 1). Addition of 1 mM (S1C1) or 5 mM (S1C5) caffeine did not significantly affect PER levels. At 7.5 mM caffeine (S1C7.5), PER decreased to 50%, suggesting partial aversion. At 10 mM caffeine (S1C10), only 20% of flies extended their proboscis, demonstrating a robust, reproducible aversive response (Fig. 1a; Video 2). Therefore, S1C10 was selected as the optimal test solution for subsequent TrpA1 experiments.

**Figure 1.**
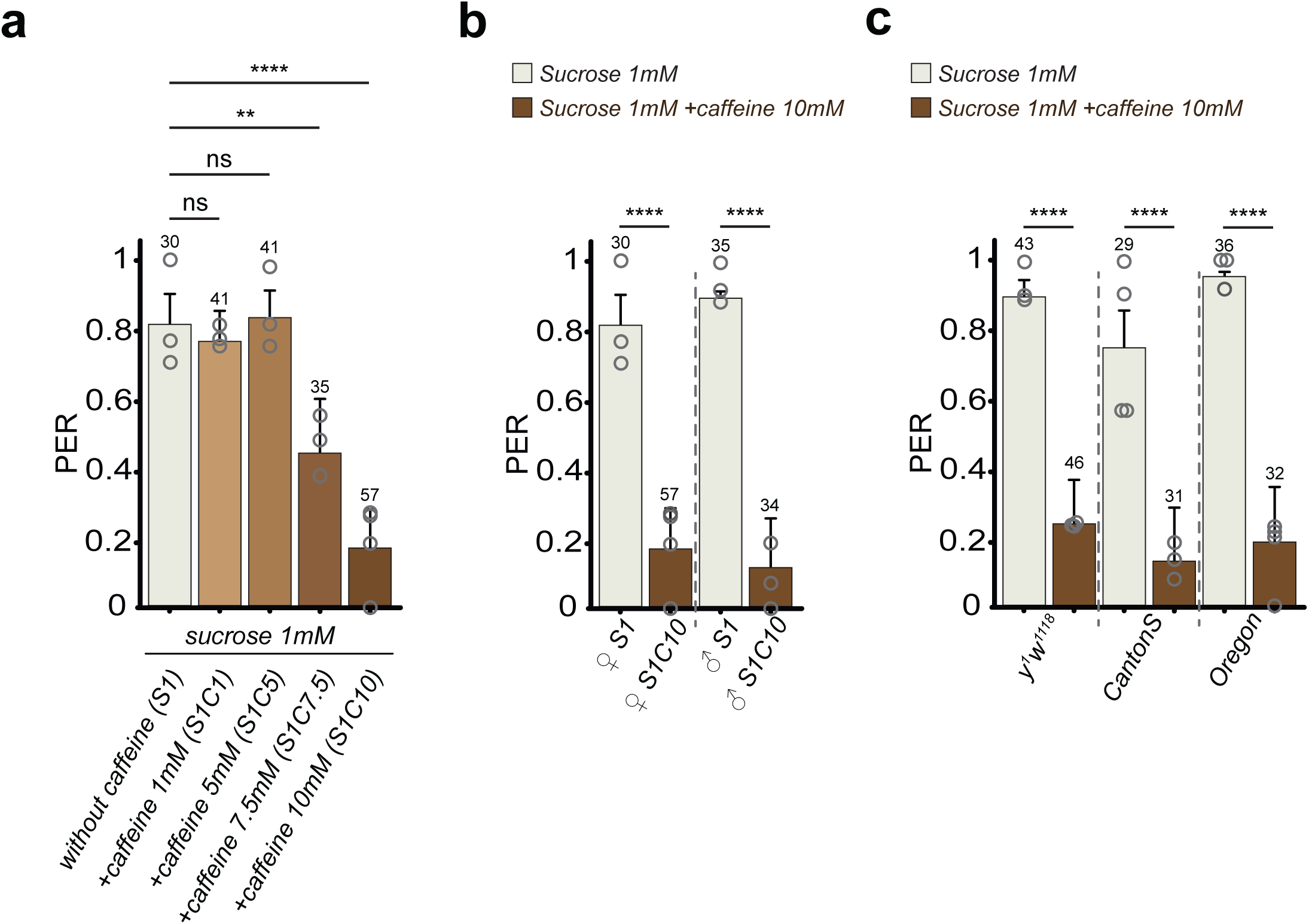
Dose-dependent caffeine inhibition of PER. (**a**) Dose dependent aversion to a mixture combining sucrose and caffeine. PER index of *w^1118^* female flies to control solutions of sucrose and sucrose + increasing concentrations of caffeine. Below the x-axis, S1 corresponds to sucrose 1mM, S1C1 corresponds to a mixture with sucrose 1mM and caffeine 1mM, S1C5 corresponds to a mixture with sucrose 1mM and caffeine 5mM, S1C7.5 corresponds to a mixture with sucrose 1mM and caffeine 7.5 mM and S1C10 corresponds to a mixture with sucrose 1mM and caffeine 10mM. (**b** and **c**) Aversion to a mixture combining sucrose and caffeine is not sexually dimorphic and independent of genetic background. PER index of females and males *w^1118^* (**b**) and *y^1^ w^1118^* or *CantonS* or *Oregon* female flies (**c**) to control solutions of sucrose and sucrose +caffeine. Labellar PER was measured to 1 mM sucrose plus or minus the indicated chemical. The PER index is calculated as the percentage of flies tested that responded with a PER to the stimulation ± 95% confidence interval (CI). A PER value of 1 means that 100% of the tested flies extended their proboscis following contact with the mixture, a value of 0,2 means that 20% of the animals extended their proboscis. The number of tested flies (n) is indicated on top of each bar. For each condition, at least 3 groups with a minimum of 10 flies per group were used. ns indicates p>0.05, ** indicates p<0.01, **** indicates p<0.0001 Fisher Exact Test. Further details can be found in the material and methods section and in the source data file.

### Flies from both sexes and of different genetic backgrounds show Caffeine-Induced Aversive Responses

Previous studies on taste perception and behavior in *Drosophila* focused primarily on female flies (Kurz et al., 2017; Masuzzo et al., 2022, 2019; Montanari et al., 2024), largely due to females’ greater resistance to starvation protocols required to increase responsiveness in PER assays. To assess potential sexual dimorphism, we measured PER responses in males exposed to S1C10. Both sexes exhibited caffeine aversion, with no significant difference between males and females (Fig. 1b). Under these conditions, sexual dimorphism in caffeine-induced PER suppression was not detected. Consequently, all subsequent experiments were conducted using 5- to 7-day-old female flies. Genetic background can affect behavioral responses in *Drosophila* (Zimmerman et al., 2012). To ensure that caffeine-induced PER suppression was not strain-specific, we tested control lines including *w⁻*, *yw*, Canton-S, and Oregon-R. All genotypes exhibited strong PER to 1 mM sucrose and significant PER reduction upon addition of 10 mM caffeine (S1C10) (Fig. 1c). These results demonstrate that caffeine-induced aversion is conserved across these genetic backgrounds.

### The TrpA1 and PLC Pathways in Neurons Are Essential for Caffeine-Induced Aversion

In *Drosophila*, taste perception has been studied extensively, and the cellular and molecular bases of avoidance behaviors are relatively well understood (Weiss et al., 2011). A central aspect of aversive taste responses is the activity of neurons expressing specific gustatory receptors (GRs). These neurons can be visualized and targeted based on their gene expression profiles. Among them, *Gr66a+* neurons are widely used as markers for bitter-sensitive cells, and Gr66a itself is essential for caffeine detection (Lee et al., 2009). Accordingly, *Gr66a+* neurons play a key role in mediating aversive feeding behaviors (Joseph and Heberlein, 2012; Moon et al., 2006) (Fig. 2a and aʹ).

**Figure 2.**
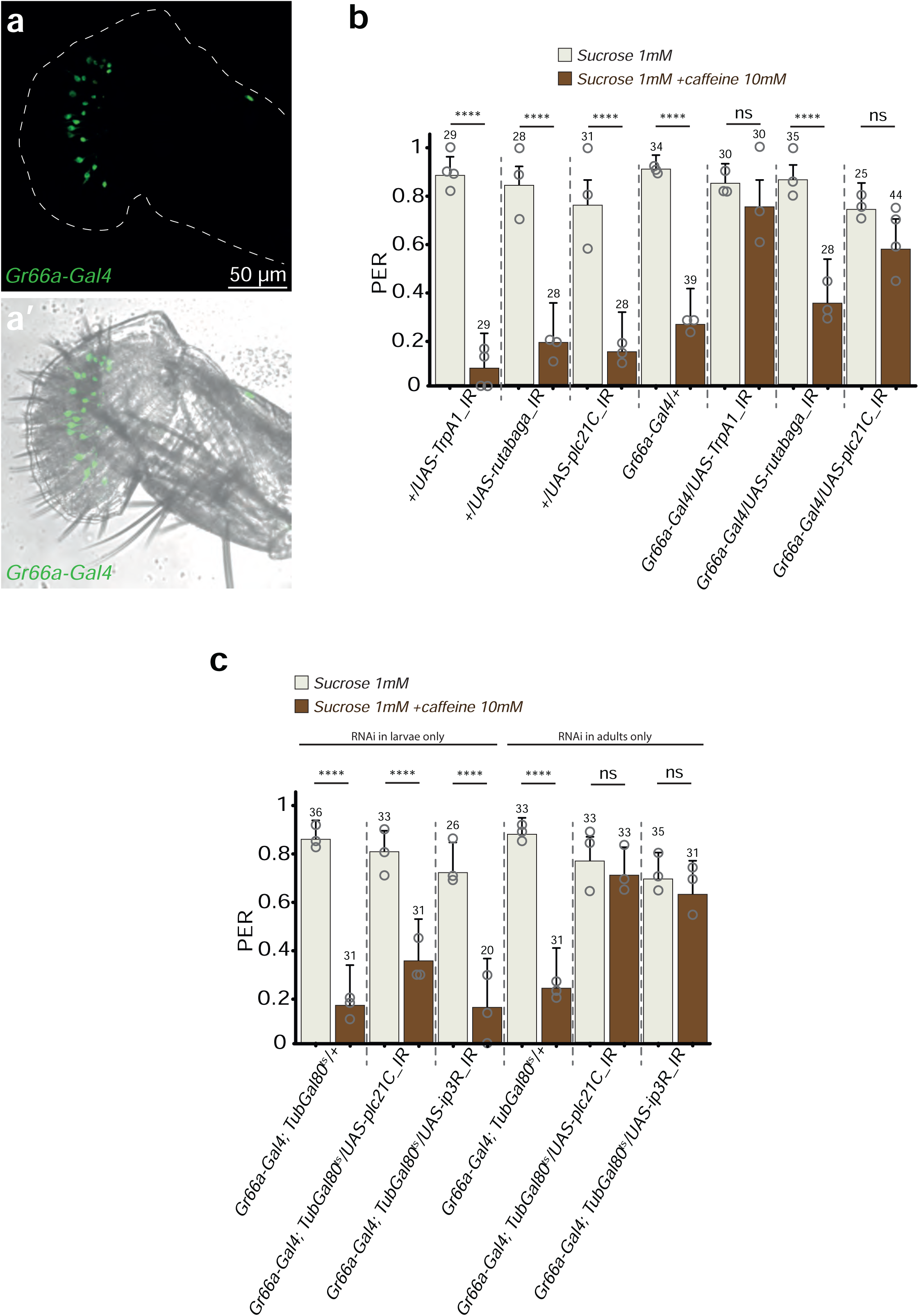
In cells the TrpA1 channel, a pathway involving Phospolipase C (PLC) and Innositol-3-phosphate receptor (IP3-R) are necessary for the caffeine-triggered aversion. (**a**) Representative confocal fluorescence image of the proboscis of *Gr66a-Gal4/UAS-GFP* flies. **(a’)** Merged image with transmitted light. (**b**) While RNAi-mediated inhibition of AMPc synthesis (*UAS-rutabaga_IR*) in cells (*Gr66a-Gal4/UAS-rutabaga_IR*) does not affect caffeine-induced aversion, RNAi-mediated inhibition of *TrpA1* (*TrpA1_IR*) in the cells (*Gr66a-Gal4/UAS-TrpA1_IR*) and of PLC21C (*UAS-plc21C_IR*) in cells (*Gr66a-Gal4/UAS-plc21C_IR*) does impair this response. PER index of flies to control solution of sucrose and to sucrose + caffeine. (**c**) PLC21C and IP3-R are functionally required in adult neurons for caffeine-triggered aversion, but are dispensable during larval life. RNAi-mediated inhibition of PLC21C (*UAS-plc21C_IR*) in cells (*Gr66a-Gal4/UAS-plc21C_IR*) and of IP3-R (*UAS-ip3R_IR*) in cells (*Gr66a-Gal4/UAS-ip3R_IR*) was controlled in a precise and timely manner, turning *on* when needed and *off* when required using *Tub-Gal80^ts^*. The ubiquitously expressed *Tub-Gal80^ts^*, that inhibits the activity of Gal4, is temperature sensitive: it’s active at 18***°***C and inactivated at 29***°***C, allowing the expression of RNAi constructions in neurons. For (**b** and **c**), PER index is calculated as the percentage of flies tested that responded with a PER to the stimulation ± 95% CI. The number of tested flies (n) is indicated on top of each bar. For each condition, at least 3 groups with a minimum of 10 flies per group were used. ns indicates p>0.05, **** indicates p<0.0001 Fisher Exact Test. Further details can be found in the material and methods section and in the source data file.

In our previous work, we confirmed that *Gr66a+* cells are necessary for caffeine-induced aversion in the PER assay. To test this, we inactivated these neurons by overexpressing the inward rectifier potassium channel Kir2.1, which impairs action potential generation throughout development (*Gr66a-Gal4/UAS-Kir2.1*) (Baines et al., 2001; Hodge, 2009; Montanari et al., 2024). As expected, this manipulation strongly reduced PER responses to S1C10, confirming that *Gr66a+* neurons are required for caffeine perception (Moon et al., 2006).

Having established their importance, we then investigated the signaling pathways within *Gr66a+* neurons that mediate caffeine aversion. Both Gr66a and Gr93a proteins have been shown to be essential for caffeine responses in preference tests, electrophysiology, and PER assays (Lee et al., 2009; Montanari et al., 2024; Moon et al., 2006). Given the involvement of the TRP channel TrpA1 in caffeine aversion in mammals (Nagatomo and Kubo, 2008), we hypothesized that it could also contribute to PER suppression in flies. The *TrpA1* gene produces five protein isoforms (A–E) through alternative splicing (Gu et al., 2019; Zhong et al., 2012). To test whether *TrpA1* in *Gr66a+* neurons is required for caffeine-induced aversion, we used a cell-specific RNA interference approach. Because the different isoforms share high sequence homology, isoform-specific RNAi was not feasible; instead, we employed a pan-isoform RNAi under the *Gr66a-Gal4 driver*. Flies carrying only the RNAi construct (without the driver) or only the driver displayed normal PER responses, extending the proboscis to sucrose and reducing it when caffeine was added. By contrast, when *TrpA1* was selectively knocked down in *Gr66a+* neurons (*Gr66a-Gal4/UAS-TrpA1_IR*), flies failed to discriminate between sucrose and the sucrose-caffeine mixture, showing no reduction in proboscis extension (Fig. 2b). These results confirm that *Gr66a+* neurons mediate the aversive response to S1C10 and demonstrate that TrpA1 activity within these neurons is essential for the immediate behavioral response to caffeine.

In the context of taste perception, intracellular signaling cascades downstream of receptor activation—particularly those involving phosphodiesterase (PDE) and phospholipase C (PLC)—have been implicated in the neuronal transduction of taste stimuli (Kwon et al., 2010; Poole and Tordoff, 2017). To assess whether these canonical pathways contribute to caffeine-induced aversion in the PER assay (S1C10), we selectively silenced *rutabaga*, which encodes PDE (Levin et al., 1992), and *plc21C* (Shortridge et al., 1991) transcripts in *Gr66a+* neurons using RNA interference. Flies were then tested for their proboscis extension reflex in response to sucrose alone or to a sucrose-caffeine mixture. Knockdown of *plc21C* in *Gr66a+* neurons markedly impaired the aversive response to sucrose + caffeine, whereas silencing *rutabaga* had no detectable effect (Fig. 2b). These findings identify PLC signaling as a critical component of the molecular cascade underlying caffeine-induced suppression of the PER in *Gr66a+* cells.

### Stage-Specific Requirement of PLC and IP3 Receptor in Caffeine Aversion

In our previous work, we demonstrated that early-life experiences can shape adult taste discrimination in *Drosophila*, particularly for molecules of bacterial origin (Montanari et al., 2024). To determine whether the role of PLC in caffeine-induced aversion is required specifically in adulthood or influenced by its absence during development, we conditionally inactivated *plc21C* in *Gr66a+* neurons at distinct life stages using the thermosensitive Gal4 repressor *Tub-Gal80^ts^*. As shown in Fig. 2c, silencing *plc21C* during the adult stage markedly impaired the aversive PER response to sucrose + caffeine, whereas its inactivation during the larval stage had no detectable effect on the adult phenotype. These findings indicate that PLC signaling in *Gr66a+* neurons is required specifically during adulthood to mediate caffeine-induced aversion. To further dissect this signaling cascade, we tested whether the inositol 1,4,5-trisphosphate receptor (IP3R) is also required in adulthood for the aversive response. Using the same conditional approach (*Gr66a-Gal4; Tub-Gal80^ts^/UAS-ip3R_IR*), we found that adult-stage inactivation of IP₃R similarly abolished the reduction in PER upon caffeine exposure (Fig. 2c). Together, these results identify PLC and IP3R as key, adult-specific components of the intracellular signaling pathway mediating caffeine aversion in *Gr66a+* neurons.

### TrpA1-E isoform mediates the response to caffeine

After establishing through cell-specific RNA interference that TrpA1 and the PLC pathway are required for caffeine aversion in *Gr66a+* neurons, we next asked which TrpA1 isoform mediates this response (Fig. 3a). Genetic tools exist in which the entire *TrpA1* gene is deleted and replaced with constructs allowing selective expression of one or more isoforms (Gu et al., 2019). These lines enable functional studies to determine whether a given isoform is necessary or sufficient for a particular behavior (Gong et al., 2023, 2022; Gu et al., 2019). Importantly, prior work using these genetic tools demonstrated that all *TrpA1+* cells are also *Gr66a+* (Sato et al., 2023). We first examined the PER in *TrpA1* null mutants lacking the entire gene. In control flies, sucrose (1 mM) induces robust proboscis extension, whereas the sucrose–caffeine mixture (S1C10) elicits aversion, reflected by suppressed extension. By contrast, *TrpA1* mutants extended their proboscis equally to both solutions (Fig. 3b), confirming that *TrpA1* is essential for caffeine-induced aversion and corroborating our RNAi results (Fig. 2b). To identify the isoform responsible for caffeine avoidance, we tested three isoform-specific knockout (KO) lines: *AD-KO*, *BC-KO*, and *E-KO* (Fig. 3a and 3b). All lines extended their proboscis to sucrose alone but only the *E-KO* line failed to reduce PER in response to the sucrose–caffeine mixture, indicating a loss of aversion (Fig. 3b). These results suggest that isoforms A–D are dispensable, whereas the E isoform is critical for caffeine-induced aversion. Notably, *E-KO* flies were tested in other studies and do not exhibit general deficits in TrpA1-dependent behaviors, such as responses to heat or reactive oxygen species (ROS) (Gu et al., 2019; Zhong et al., 2012), suggesting a specific role for the E isoform in bitter taste signaling.

**Figure 3.**
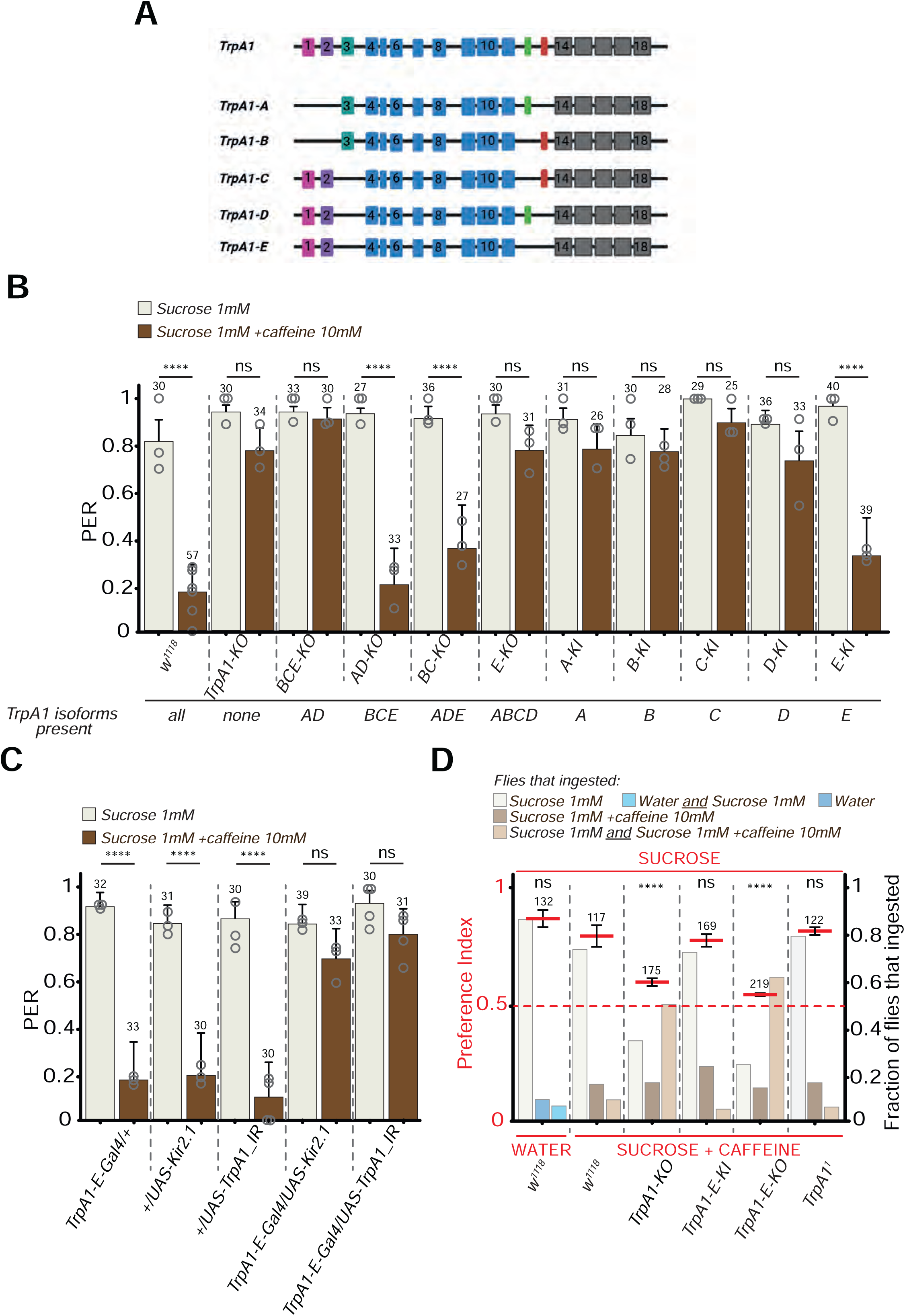
The TrpA1-E-isoform and the *TrpA1-E+* cells are necessary for the caffeine-triggered aversion. (**a**) Graphical representation of the characterized TrpA1 isoforms produced by the *TrpA1* gene.A/B/C/D/E isoforms share the blue and grey exons, C/D/E share the exons 1 and 2, A and B share the exon 3, A and D share the exon 12, B and C share the exon 13. E is the only isoform not having exon 12 or exon 13. On top is a representation of the *TrpA1* gene with all the exons. (**b**) The TrpA1-E isoform is necessary for the caffeine-triggered aversion in adult flies. Mutant lines impaired for the production of all the isoforms (*TrpA1-KO*) or producing only one specific isoforms (-KI) were tested. For every line assayed, below the x-axis, the TrpA1 isoforms still produced are indicated. (**c**) Impairing the activity of *TrpA1-E+* cell via *UAS-Kir2.1* (*TrpA1-E-Gal4/UAS-Kir2.1*) abrogates caffeine-triggered aversion and RNAi-mediated *TrpA1* inactivation (*UAS-TrpA1_IR*) in the *TrpA1-E+* cells (*TrpA1-E-Gal4/UAS-TrpA1_IR*) abrogates the caffeine-induced aversion. (**d**) The TrpA1-E isoform is necessary for the caffeine-triggered aversion in a 2-choices assay. Shown are the preference indexes for different mutants as well as the fraction of flies that ingested the proposed mixtures. For (**b** and **c**) PER index is calculated as the percentage of flies tested that responded with a PER to the stimulation ± 95% CI. The number of tested flies (n) is indicated on top of each bar. ns indicates p>0.05, **** indicates p<0.0001 Fisher Exact Test. Further details can be found in the material and methods section and in the source data file. For (**d**), the averaged preference index is in red with the standard error of the mean obtained from 6 to 8 independent experiments. The total amount of females tested is indicated on top of each assay. A preference index (PI) over 0.5 indicates a preference toward sucrose while a PI of 0.5 indicates either that two opposite behaviors occurred in the population or that a majority of animals consumed both solutions. The fraction of flies that ingested one, the other or both solutions allow to discriminate. In light colors are the fractions of flies that ingested one solution or both solutions. The animals from all the experiments were pooled for statistical analyses. ns indicates p>0.05, **** indicates p<0.0001 Chi-square Test. Further details can be found in the material and methods section and in the source data file.

To further validate this finding, we tested knock-in (*KI*) lines that can only express one specific TrpA1 isoform (A–E) (Gu et al., 2019). Flies expressing isoforms A–D displayed PER responses similar to *TrpA1* null mutants, showing no aversion to sucrose–caffeine mixtures. In contrast, the *E-KI* line behaved like wild-type flies, suppressing PER in response to caffeine, confirming that the E isoform is necessary for this behavior (Fig. 3b). Finally, crossing the *A-KI* and *D-KI* lines to generate *BCE-KO* flies, which lack isoforms B, C, and E, recapitulated the *E-KO* phenotype, further supporting the specific requirement of TrpA1-E for caffeine avoidance.

Taken together, these data demonstrate that the TrpA1-E isoform is necessary for PER suppression when caffeine is added to a sucrose solution. To determine whether the TrpA1-E isoform was necessary for aversion to other chemicals, we tested the PER responses of *TrpA1-E-KO* and *TrpA1-E-KI* flies to quinine. The results demonstrate that the TrpA1-E isoform is required for aversion to quinine, as well as to caffeine (Supplementary Fig. S1). As all *TrpA1+* cells also express *Gr66a* (Sato et al., 2023), this finding supports a central role for TrpA1-E in *Gr66a+* cells in mediating avoidance behavior toward aversive compounds.

The requirement of the TrpA1-E isoform for caffeine-induced aversion suggests that the cells expressing this isoform are directly responsible for the observed behavior. To test this, we performed PER assays in flies in which *TrpA1-E+* cells were silenced by overexpressing the inward-rectifier potassium channel Kir2.1 throughout development (larvae, pupae, and adults), effectively impairing action potential generation (*TrpA1-E-Gal4/UAS-Kir2.1*). Control flies carrying the UAS transgene without the driver *(+/UAS-Kir2.1*) or only the driver (*TrpA1-E-Gal4/+*) extended their proboscis in response to sucrose but not to the sucrose–caffeine mixture (S1C10), consistent with expected aversion (Fig. 3c). In contrast, flies with silenced *TrpA1-E+* cells responded normally to sucrose but failed to suppress PER when caffeine was added, indicating that these neurons are necessary for the reduction of proboscis extension in the presence of caffeine (Fig. 3c).

To confirm that this effect depends specifically on TrpA1 activity in *TrpA1-E+* cells, we used a pan-isoform RNAi construct under the control of the *TrpA1-E-Gal4* driver. Flies expressing only the RNAi transgene *(+/UAS-TrpA1_IR*) behaved like controls, extending their proboscis to sucrose and reducing PER in response to sucrose–caffeine. By contrast, *TrpA1-E+ cells-*specific RNAi (*TrpA1-E-Gal4/UAS-TrpA1_IR*) completely abolished caffeine avoidance, phenocopying the *TrpA1* null mutants (Fig. 3b and 3c).

To further substantiate the role of *TrpA1-E+* cells in caffeine aversion, we performed two-choice feeding assays using 96-well plates with colored agar overlaid with test solutions (Weiss et al., 2011). Freely moving females were allowed to feed, and abdominal coloration was used to assess substrate preference (Supplementary Fig. S2). Control (*w⁻*) flies strongly preferred 1 mM sucrose over water and over sucrose–caffeine mixtures (preference index > 0.79) (Fig. 3d). In contrast, *TrpA1-KO* and *TrpA1-E-KO* flies showed markedly reduced avoidance of the sucrose–caffeine solution (preference index < 0.6), with most individuals consuming both solutions (Fig. 3d). *TrpA1-E-KI* flies behaved like controls, demonstrating robust avoidance (preference index > 0.75). Interestingly, the *TrpA1¹* allele used in the pioneer study (Kim et al., 2010) also responded similarly to wild-type flies, suggesting that this mutation may not completely abolish TrpA1 function or may differentially affect specific isoforms (Fig. 3d). Together, these findings provide compelling evidence that *TrpA1-E+* cells are required for caffeine aversion, likely mediated specifically by the TrpA1-E isoform. Furthermore, results from cell-targeted RNAi and multiple mutants, including *TrpA1¹*, suggest that the *TrpA1¹* mutation may not produce a complete loss of function or may differentially affect TrpA1 isoforms.

### *TrpA1-E+* cells are taste neurons of the proboscis that respond to caffeine

Having established that *TrpA1-E+* cells are necessary for the PER response to the sucrose–caffeine mixture (S1C10), and noting that previous work using a pan-isoform *TrpA1* driver showed that all *TrpA1+* cells are also *Gr66a+* (Sato et al., 2023), we next investigated the specific localization of *TrpA1-E+* cells within the proboscis, focusing on their association with taste sensilla. Such neurons have previously been implicated in aversive behaviors triggered by chemical contact with proboscis taste sensilla (Liman et al., 2014) To investigate this, we imaged the proboscis of female flies expressing GFP under the control of the *TrpA1-E-Gal4* driver. On average, we observed ∼10 *TrpA1-E+* cells per labellum, each sending a distinct projection backward into the pharynx. Additional processes extended into the taste sensilla, reaching their tips (Fig. 4a–4aʺ). These morphological features are consistent with previously described proboscis taste neurons (Liman et al., 2014). Because the PER is a rapid reflex triggered by contact of the proboscis taste sensilla with a stimulus-soaked surface, our anatomical and functional data strongly suggest that the reduction of PER in response to sucrose–caffeine (S1C10) is mediated by *TrpA1-E* expressed in these specific proboscis taste neurons. Thus, *TrpA1-E+* cells within the proboscis are integral to caffeine-induced aversion.

**Figure 4.**
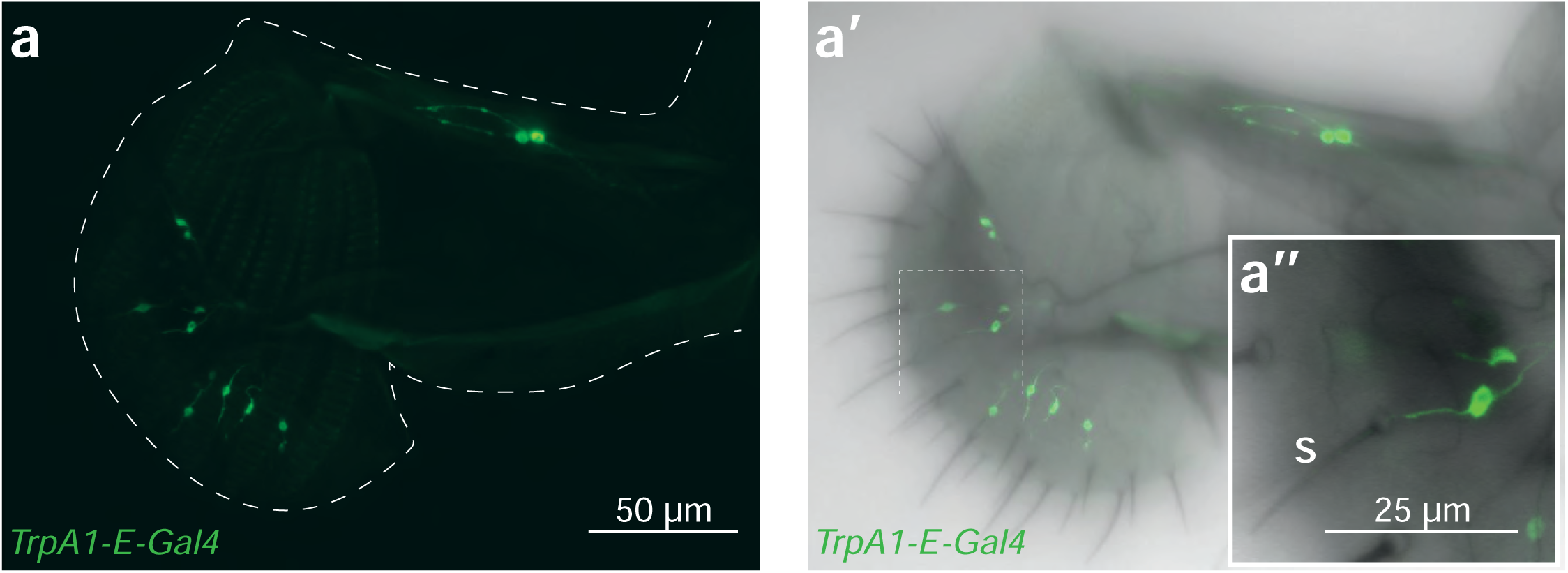
The *TrpA1-E+* cells in the labellum have the characteristics of peripheral taste neurons. **(a)** Representative confocal fluorescence image of the proboscis of *TrpA1-E-Gal4/UAS-GFP* flies. **(a’)** Merged image with transmitted light. **(a”)** Magnified view of the area delineated by the white dashed line, showing a cell with an extension inserting into a short sensillum (S) and another extension running posteriorly.

To further confirm that *TrpA1-E+* cells directly respond to caffeine, we monitored intracellular calcium dynamics using GCaMP (Jayaraman and Laurent, 2007). Proboscises were exposed to caffeine, and fluorescence changes in the cell bodies of *TrpA1-E+* neurons were quantified (*TrpA1-E-Gal4/UAS-GCaMP*). These experiments revealed that *TrpA1-E+* cells respond directly to caffeine (Fig. 5). Importantly, this calcium response was abolished when *TrpA1* was knocked down specifically in *TrpA1-E+* cells using RNAi (*TrpA1-E-Gal4/UAS-GCaMP; UAS-TrpA1_IR*) (Fig. 5), confirming that TrpA1 is required for the direct detection of caffeine by these neurons.

**Figure 5.**
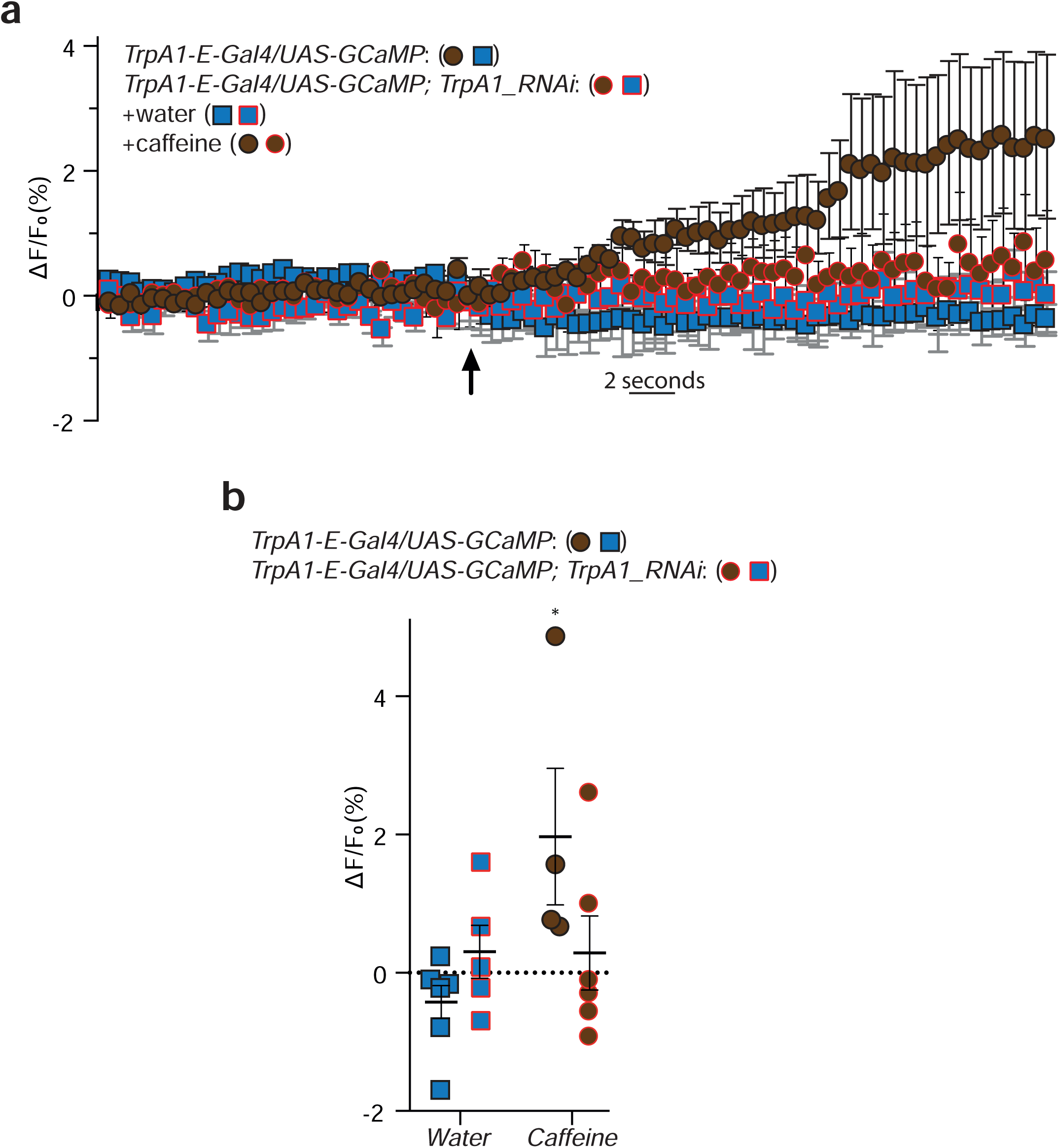
*TrpA1-E+* cells located in the proboscis respond to caffeine exposure. **(a)** Averaged ± SEM time course of the GCaMP intensity variations (ΔF/F0 %) for proboscis *TrpA1-E+* cells of *TrpA1-E-Gal4/UAS-GCaMP* and *TrpA1-E-Gal4/UAS-GCaMP; UAS-TrpA1_IR* females. The addition of water (n = 5 and 5 flies) or caffeine 10mM (n = 4 and 6 flies) at a specific time is indicated by the arrow. Several neurons in the focal plan located in the same proboscis were used for the quantification. **(b)** Averaged fluorescence intensity of peaks ± SEM in response to water (n = 5 and 5 flies), caffeine 10mM (n = 4 and 6 flies). *p =0.046, Kruskal-Wallis test compairing the 4 independent sets of values.

## DISCUSSION

Previous studies have examined the effects of caffeine on adult flies using various approaches, including the CAFé assay (Sellier et al., 2011), two-choice assays (Kim et al., 2010), and electrophysiological recordings (Kim et al., 2010). These studies identified key receptors, such as Gr66a (Moon et al., 2006) and Gr93a (Lee et al., 2009), as essential for caffeine perception. TrpA1, a well-characterized sensor in flies, has also been implicated in aversion to certain chemicals, including aristolochic acid (Kang et al., 2010; Kim et al., 2010; Kwon et al., 2010). Interestingly, when assayed in some of these studies, the *TrpA1* mutants used still exhibited caffeine avoidance in behavioral assays and electrophysiological responses, leading to the conclusion that TrpA1 might not be required for caffeine aversion in adult flies (Kim et al., 2010). In contrast, our findings show that TrpA1 is indeed necessary for caffeine avoidance in PER and two-choice assays, as well as for *TrpA1+* cell activity following caffeine exposure, as demonstrated by our experiments using knockout (KO), knock-in (KI) mutants, RNAi and GCaMP.

The primary assay in our study was the proboscis extension reflex (PER), which differs from two-choice assays. In a typical two-choice assay, food-deprived flies select between two solutions, one containing caffeine. The PER assay, however, measures a fly’s immediate reflex upon contact with a substance, providing a more direct readout of taste response. Notably, caffeine (10 mM) mixed with 5 mM sucrose causes aversion in two-choice assays, whereas in the PER assay, caffeine mixed with sucrose concentrations above 2 mM is no longer aversive (Montanari et al., 2024). While two-choice assays capture physiologically relevant, group-based decision-making over a longer period and can involve additional sensory inputs such as leg-mediated taste or odor, the PER reflects rapid, reflexive responses. In our tests, *TrpA1* mutants failed to respond to caffeine in two-choice assays, whereas the *TrpA1^1^* allele did, suggesting that this allele may not completely disrupt all TrpA1 functions or isoforms.

Electrophysiological measurements, which assess immediate neuronal responses, also differ from behavioral PER outcomes. *TrpA1* is expressed in several taste neurons, including those of the S6 sensilla (Kwon et al., 2010). Bitter neurons in these sensilla may respond to caffeine independently of TrpA1, as observed with citronellal, which triggers both TrpA1-dependent and -independent responses (Kwon et al., 2010). The discovery of five TrpA1 isoforms with distinct functions further complicates the interpretation of activation patterns. Genetic compensation is unlikely to explain discrepancies, as our mutants and RNAi treatments produced consistent phenotypes (Dy et al., 2024). Therefore, differences between our results and prior studies likely reflect variations in assay type, sensilla tested, and the specific effects of mutations on the newly described TrpA1 isoforms.

### TrpA1 and PLC/IP3R signaling

Our study supports a conserved relationship between TrpA1 and PLC signaling observed in both mammalian (Zhang et al., 2003) and *Drosophila* models (Kim et al., 2010; Kwon et al., 2010). Gustatory responses to sweet and bitter stimuli often involve TRP channels, such as TRPM5, and phospholipase C (PLCβ2) (Zhang et al., 2003). In insects, TrpA1 functions downstream of PLC (Kwon et al., 2008). PLC activity generates inositol trisphosphate (IP3) and diacylglycerol (DAG); DAG may directly activate TrpA1 to allow ion influx, while IP3 releases calcium from ER stores via the IP3 receptor (IP3R), altering cytoplasmic calcium levels and triggering action potentials. In taste neurons, this cascade likely involves upstream elements, including G-alpha proteins and caffeine-activated GPCRs, and enhances signal specificity. TrpA1 acts as a molecular integrator, converting PLC-induced Ca²⁺ fluxes into depolarizing currents with temporally regulated dynamics. Rapid IP3-mediated Ca²⁺ spikes may drive acute aversion, while sustained DAG signaling could mediate adaptation.

### The E-isoform of TrpA1

First described in 2019, the E-isoform of TrpA1 had not been assigned a specific function, unlike the B- and C-isoforms (Gu et al., 2019). Our data show that TrpA1-E is essential for the reflexive response to caffeine. While it may not act as a direct caffeine sensor, TrpA1 channels can function as “alarm signals,” depolarizing neurons following receptor-ligand interactions. Thus, both the expression of TrpA1 and its interaction with gustatory receptors are crucial. The functional distinctions of each isoform likely dictate these interactions. Notably, TrpA1-E is required for PER aversion to caffeine and is expressed in fewer than ten neurons, compared to approximately thirty *Gr66a+* cells per labellum involved in this response, suggesting that the E-isoform defines a critical subset of *Gr66a+* neurons essential for caffeine detection.

## Acknowledgments

This work was supported by CNRS, ANR BACNEURODRO (ANR-17-CE16-0023-01), Equipe Fondation pour la Recherche Médicale (EQU201603007783) et l’Institut Universitaire de France to J.R. and the ANR Pepneuron (ANR-21-CE16-0027) to J.R. and Y.G. Research in Y.G.’s laboratory is supported by the CNRS, the “Université de Bourgogne Europe”, the Conseil Régional Bourgogne Franche-Comté (PARI grant), the FEDER (European Funding for Regional Economical Development), and the European Council (ERC starting grant, GliSFCo-311403).

## MATERIAL AND METHODS

### Fly husbandry

Flies were grown at 25°C on a yeast/cornmeal medium in 12h/12h light/dark cycle-controlled incubators. For 1 L of food, 8.2 g of agar (VWR, cat. #20768.361), 80 g of cornmeal flour (Westhove, Farigel maize H1) and 80 g of yeast extract (VWR, cat. #24979.413) were cooked for 10 min in boiling water. 5.2 g of Methylparaben sodium salt (MERCK, cat. #106756) and 4 mL of 99% propionic acid (CARLOERBA, cat. #409553) were added when the food had cooled down. We use a protein-rich rearing media, in contrast to the sugar-rich media used in some other laboratories. Consequently, our animals may be more sensitive to low sucrose concentrations (1 mM) compared to flies raised on sugar-rich media. Therefore, assays using 1 mM sucrose may not yield reproducible results when performed with flies reared on sugar-rich media.

### Fly stocks

The reference strain in this study corresponds to *w^[1118]^* (Bloomington #5905). Other lines were Canton S (Bloomington #64349) and *y^[1]^, w^[1118]^* (gift from Bernard Charroux) and *Gr66a-Gal4* (Bloomington #28801) and *UAS-Kir2.1* (Bloomington #6595) and *Tub-Gal80^ts^* (Bloomington #7016) and *UAS-Rutabaga_IR* (Bloomington #27035) and *UAS-GFP* (Bloomington #32195) and *UAS-PLC21C_IR* (Bloomington #31270) and *UAS-TrpA1_IR* (Bloomington #36780) and UAS-*IP3R_IR* (Bloomington #25937) and *TrpA1^1^* (Bloomington #26504) and UAS-GCaMP7s (Gift from Matthieu Cavey). The *UAS-TrpA1_IR* (Bloomington #36780) should target all the isoforms as it corresponds to a sequence in exon 17, an exon shared by all the isoforms. All the *TrpA1* isoform-specific genetic tools (*TrpA1-KO* and *TrpA1-AD-KO* and *TrpA1-BC-KO* and *TrpA1-E-KO* and *TrpA1-A-KI* and *TrpA-1-B-KI* and *TrpA1-C-KI* and *TrpA1-D-KI* and *TrpA1-E-KI* and *TrpA1-E-Gal4*) were kindly provided by the Yang Xiang laboratory. The *TrpA1-BCE-KO* is an F1 obtained after having crossed together the *TrpA1-A-KI* with the *TrpA1-D-KI*. These *TrpA1* genetic tool kit lines are based on a complete *TrpA1* gene deletion, replaced in situ with constructs allowing the specific expression of one or several isoforms, including *Gal4* driver constructs (Gu et al., 2019). These fly lines make it possible, for example, to investigate whether a given isoform is necessary or sufficient for a particular mechanism (Gong et al., 2023).

### RNAi timely controlled

The ubiquitously expressed *Tub-Gal80^ts^*, that inhibits the activity of Gal4, is temperature sensitive: it’s active at 18**°**C and inactivated at 29**°**C, allowing the expression of *UAS* when animals are raised at 29**°**C and preventing it at 18**°**C. For RNAi in larvae only, animals were raised from eggs to early pupae at 29**°**C and then shifted at 18**°**C. for RNAi in adults only, eggs, larvae and pupae were raised at 18**°**C and the virgin adults shifted to 29°C.

### PER assay

All flies used for the test were females between 5 and 7 days old. Unless experimental conditions require it, the flies are kept and staged at 25°C to avoid any temperature changes once they are put into starvation. The day before, the tested flies are starved in an empty tube with water-soaked plug for 24h at 25°C.

Eighteen flies are tested in one assay, 6 flies are mounted on one slide and in pairs under each coverslip. To prepare the slide, three pieces of double-sided tape are regularly spaced on a slide. Two spacers are created on the sides of each piece of tape by shaping two thin cylinders of UHT paste. To avoid the use of carbon dioxide flies are anesthetized on ice. Under the microscope, two flies are stuck on their backs, side by side, on same piece of tape so that their wings adhere to the tape. A coverslip is then placed on top of the two flies and pressed onto the UHT paste, blocking their front legs and immobilizing them.

Once all slides are prepared, they are transferred to a humid chamber and kept at 25°C for 1.5 hours to allow the flies to recover before the assay.

Flies are tested in pairs, the test is carried out until completion on a pair of flies (under the same coverslip), and then move on to the next pair.

Before the test, water is given to each pair of flies to ensure that the flies are not thirsty and do not respond with a PER to the water in which the solutions are prepared. Stimulation with the test solution is always preceded and followed by a control stimulation with a sweet solution, to assess the fly’s condition and its suitability for the test. During the test small strips of filter paper are soaked in the test solution and used to contact the fly’s labellum (three consecutive times per control and test phase). Contact with the fly’s proboscis should be as gentle as possible. Ideally the head should not move. A stronger touch may prevent the fly from responding to subsequent stimulation. Based on the protocol needed the test is done following the sequence and the timing in the table below.

**Tab. 1.**
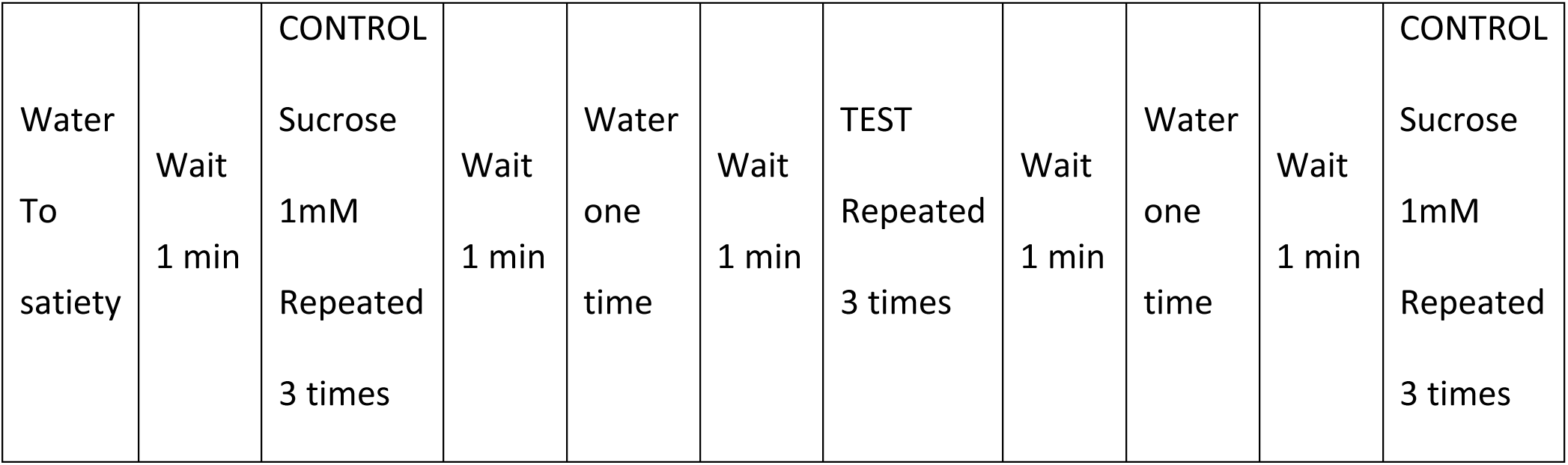
Aversion protocol.

All solutions to be tested are prepared the test day and stored at room temperature. In the aversion protocol, the control stimulation is performed with 1mM sucrose (D(+)-sucrose ≥99.5 %, p.a. Carl Roth GmbH + Co. KG). This concentration is sufficient to elicit a PER but is not so high as to influence the response to the subsequent test stimulation. After each control or test stimulation, a water-soaked strip is used to tap the proboscis and clean it. The caffeine mixed with the sucrose is Caffeine powder ReagentPlus ref C0750 Sigma Aldrich and quinine Q1125 Sigma Aldrich.

The response of the fly to each stimulation is recorded and averaged. We distinguish between two types of flies: those able to respond and those unable to respond. To make this distinction, we use three control stimulations before the test and three after the test. Flies that do not respond during the control stimulations (1 mM sucrose) are classified as “unable to respond” and are excluded from the analysis, regardless of their responses during the tests. Flies that respond during the control stimulations are classified as “able to respond,” and only their responses during the tests are included in the statistical analyses. The PER index is calculated as the percentage of flies “able to respond” tested that responded with a PER to the TEST stimulation and represented as ± 95% CI.

### Choice assay

Two-way choice assays were performed essentially as described (Meunier et al., 2003). Briefly, 5- to 7-day-old female flies (30 ± 10 flies per experiment) were starved for ∼18 h at 25°C and placed into a 96-well plate with 1% agarose in each well. Alternating wells contained either red (sulforhodamine B, 0.2 mg/mL; Sigma-Aldrich) or blue dye (bleu E133 meilleur du chef, 0.3%). Then we deposit 10µL of solution containing sucrose 1mM or Sucrose1mM + Caffein 10mM or water on top of the well. Each experiment is carried out in duplicate by placing the components on two different colors (*e.g.*, sucrose 1mM on blue wells for plate A and on red wells for plate B) to verify whether there is an effect of color. The flies were allowed to feed in the dark for 90 min at ∼ 23°C. We transferred the plates to -20°C for 15min in order to rapidly freeze the flies in order to count the flies based on their abdominal color under a binocular microscope. The numbers of flies with blue, red, or purple (mixed red and blue) abdomens were tabulated (Supplementary Fig. S2), and the P.I. was determined: (N_suc_ + 0.5N_mix_)/(N_suc_ + N_test_ + N_mix_). The flies without colors in their abdomen were discarded. The number of flies indicated corresponds to animals with colored abdomens. With *Trpa1^1^* allele, the proportion of uncolored flies was remarkably higher that with the other genotypes.

### Microscopy

No immunostaining was performed with proboscises. Proboscises of adult females were dissected in PBS, rinsed with PBS, and directly mounted on slides using Vectashield fluorescent mounting medium. The tissues were visualized directly after. Images were captured with a LSM 780 Zeiss confocal microscope (20X air objective was used).

### Calcium imaging

*In vivo* adult calcium imaging experiments were performed on proboscises of 5-7 day-old starved mated females. Animals were raised on conventional media with males at 25°C. Flies were starved for 20-24 h in a tube containing a filter paper soaked in water prior to experiments. Flies of the appropriate genotype were anesthetized on ice for 1 h. The heads of the animals were gently harvested and immobilized using insect pins (0.1 mm diameter) placed on the head and on the proboscis. 200 µL of water was placed on the head and proboscis. GCaMP7s fluorescence was viewed with a Leica DM600B microscope under a 25x water objective. Stimulation was performed manually using a pipette with gel loading tip by applying 200 µL of water or 200 µL of Caffeine (20mM of caffeine diluted in water for a final caffeine concentration on the preparation of 10mM). The gustatory stimulation was continuous as the tastant is in the solution contacting the proboscis. The recording started before the addition of the tastant and the calcium response could be observed following the contact with the proboscis. GCaMP7s was excited using a Lumencor diode light source at 482 nm ± 25. Emitted light was collected through a 505-530 nm band-pass filter. Images were collected every 500ms. Each experiment consist of the recording of 30 images before stimulation to determine a F0 (mean of the 30th images previous stimulation). Then a fluorescence variation in a Region Of Interest (ROI) is calculated as (Fi-F0)/F0 *100 from F1 to F131 and substrated to the fluorescence of another ROI that represents the background from F1 to F131. Data were analyzed as previously described (Silbering et al., 2012) by using FIJI (https://fiji.sc/).

### Statistics and data representation

The Prism software was used for statistical analyses.

For PER datasets. As the values obtained from one fly are categorical data with a *Yes* or *No* value, we used the Fisher exact t-test and the 95% confidence interval to test the statistical significance of a possible difference between a test sample and the related control.

For PER assays, at least 3 independent experiments were performed. The results from all the experiments were gathered and the total amount of flies tested is indicated in the graph. In addition, we do not show the average response from one experiment representative of the different biological replicates, but an average from all the data generated during the independent experiments in one graph. However, each open circle represents the average PER of 1 experiment.

For choice assays datasets. As the values obtained from one fly are categorical data with a *Red* or *Blue* or *Mixed* value, we used the Chi-square test to take the number of tested flies into account. At least 6 independent experiments were performed and the preference index shown is an average from the PI of each independent experiment.

For in vivo calcium imaging, the D’Agostino–Pearson test to assay whether the values are distributed normally was applied. As not all the data sets were considered normal, non-parametric statistical analysis such as Kruskal–Wallis H test was used for all the data presented. For a given genotype, the reference for the comparisons is the exposure to water.

**Detailed lines, conditions and statistics for the figure section**

All these data are downloadable with the source data file.

Private link: https://figshare.com/s/31cce8c3f518b30956c9

## VIDEOS

**Video 1 PER is triggered following exposure to sucrose 1mM**

Private link: https://figshare.com/s/d5dc0d81a2a1a3b19365

Response following proboscis contact with a stripe of paper soaked in a 1mM solution (water as solvent). 7-days old females starved 15h prior to the assay.

**Video 2 PER is not triggered following exposure to a mixture combining sucrose 1mM and caffeine 10mM**

Private link: https://figshare.com/s/e5ce4ca699699046e076

Absence of response following proboscis contact with a stripe of paper soaked in sucrose 1mM combined with caffeine 10mM. 7-days old females starved 15h prior to the assay.

## Ethics statements

N/A

## Conflict of interest statement

No conflict of interest

## Funding statement

This work was supported by CNRS, ANR Pepneuron (ANR-21-CE16-0027) to J.R. and Y.G. Research in Y.G.’s laboratory is supported by the CNRS, the “Université de Bourgogne Europe”, the Conseil Régional Bourgogne Franche-Comté (PARI grant), the FEDER (European Funding for Regional Economical Development), and the European Council (ERC starting grant, GliSFCo-311403).

## Data avaibility

All these data are downloadable with the source data file. Private link: https://figshare.com/s/31cce8c3f518b30956c9

---

**Supplementary Figure 1.**
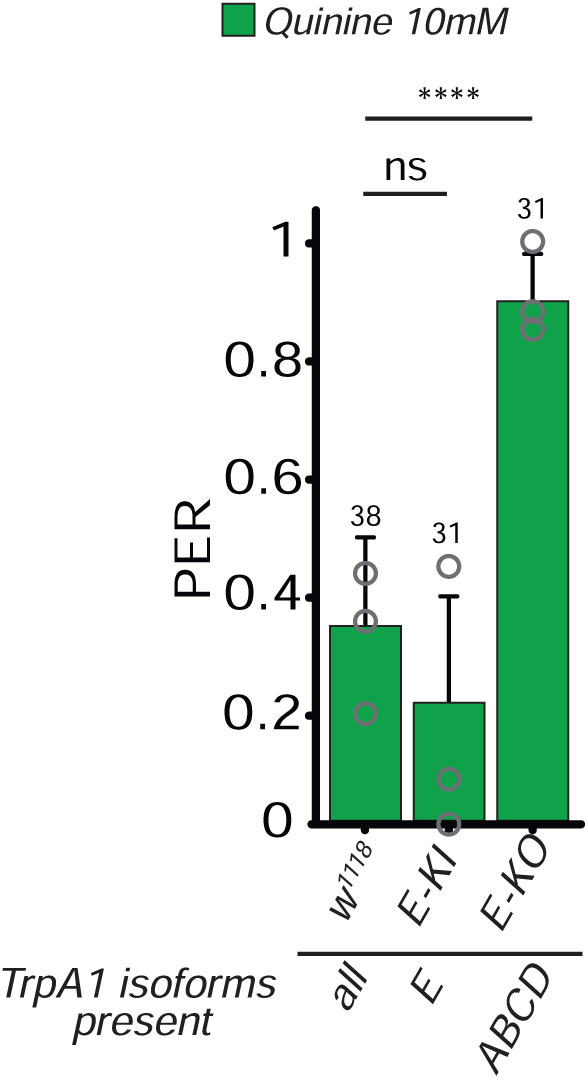
The TrpA1-E isoform is necessary for the caffeine-triggered aversion in adult flies. Control (*w1118*) and mutant lines impaired for the production of the E-isoform (*E-KO*) or producing only the E-isoform (E-KI) were tested. For every line assayed, below the x-axis, the TrpA1 isoforms still produced are indicated. PER index is calculated as the percentage of flies tested that responded with a PER to the stimulation ± 95% CI. The number of tested flies (n) is indicated on top of each bar. ns indicates p>0.05, **** indicates p<0.0001 Fisher Exact Test. Further details can be found in the material and methods section and in the source data file.

**Supplementary Figure 2.**
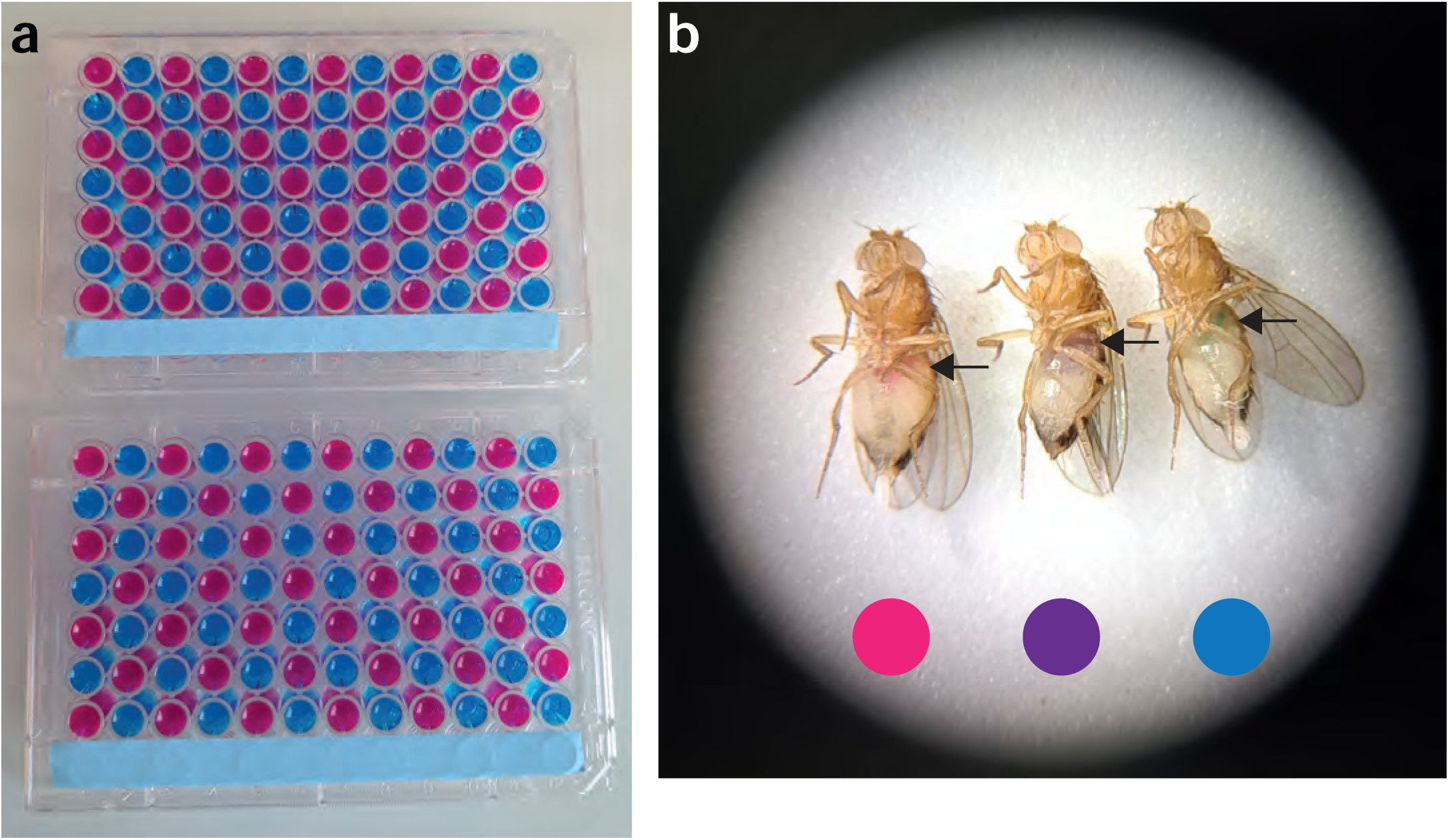
**(a)** Two-way choice assays were performed into a 96-well plate with 1% agarose in each well. Alternating wells contained either red or blue dye with 10µL of solution containing sucrose 1mM or Sucrose1mM + Caffein 10mM or water on top of the well. **(b)** Flies are then counted based on their abdominal color (arrow pointing the crop) under a binocular microscope.

## Notes

https://figshare.com/s/31cce8c3f518b30956c9

https://figshare.com/s/d5dc0d81a2a1a3b19365

https://figshare.com/s/e5ce4ca699699046e076

### Competing Interest Statement

The authors have declared no competing interest.

